# Forward Genetics Identifies *ptr-18* and Other Genes as Developmental Regulators of the Four-Cell Tail Tip in *C. elegans*

**DOI:** 10.1101/2025.08.20.671392

**Authors:** Uroš Radović, Marcus Henricsson, Jan Borén, Marc Pilon

## Abstract

In *C. elegans*, the epidermis and its overlying extracellular matrix form a primary protective barrier, functioning as the first line of defense against environmental factors. To properly develop those cellular boundaries, a tightly controlled interaction of many molecules and pathways is needed. Mutant alleles of *paqr-2* and *iglr-2* (lipid homeostasis), *dpy-21* (membrane trafficking), *and sma-1* (actin-binding spectrin) result in hermaphrodite tail tip defects suggesting that this simple four-cell structure can serve as a sensitive model for the identification of pathways responsible for the establishment of cellular boundaries. With this in mind, we performed a small forward genetics screen of ∼800 ethyl methanesulfonate-mutagenized haploid genomes and identified 21 mutants with a tail end defective (Ted) phenotype. Whole genome sequencing of these mutants identified mutations in genes encoding either structural constituents of the cuticle itself (mostly collagen genes) or protein with regulatory functions. By using CRISPR/Cas9 we confirmed six novel alleles of *ptr-18, paqr-2, nab-1, ncam-1, vab-9 and efn-4.* We further characterized the loss of function allele *ptr-18(et70)*, which encodes a patch domain-containing (PTCHD) protein homologous to human PTCHD1. *ptr-18(et70)* has a significant effect on growth and development of the worms, while also increasing membrane permeability. Lipidomics analysis revealed no major alterations in membrane lipid composition, implicating cuticle defects as the primary cause of the observed permeability phenotype.

**Article summary:** We performed a forward genetics screen to identify hermaphrodite *C. elegans* mutants with a tail end defect with the goal to discover membrane and morphogenesis regulators. The screen of 800 haploid genomes revealed 21 tail end defective mutants, including 8 novel alleles of interesting regulator protein. We conclude that the tail tip phenotype can be useful in discovery of new pathways and interactions during development.

## Introduction

Hypodermal cells in the nematode *C. elegans* form the outer cellular layer of the body and secrete a collagenous extracellular matrix (ECM) that organizes itself into cortical and basal layers separated by struts, forming the worm’s cuticle [1]. During larval development, worms replace their cuticle before each of the four molting cycle, a process regulated by the differential expression of many collagen genes [2]. Defects in cuticle composition and genetic regulation can cause a wide range of body shape phenotypes. Indeed, the genetic basis of morphology in *C. elegans* has been extensively researched by studying mutants with body shape phenotypes such as Long (Lon), Dumpy (Dpy), Small (Sma) and Variable abnormal (Vab), and many of the causative alleles affect the synthesis or trafficking of cuticle components [3]. For example, of over 30 characterized Lon and Dpy mutants, 11 affect collagen genes (there are ∼177 collagen genes in the *C. elegans* genome)[3–6].

While gross body shape mutants have been extensively characterized, there is still merit in investigating morphology mutants affecting more subtle or specific structures. For example, mutants that affect the morphology and development of the complex fan-like male tail have provided useful insight into the mechanisms of cell fate decisions and cell fusion [7, 8]. Another structure that merits careful investigation is the long and tapered hermaphrodite tail (**Fig. 1a**), which is comprised of four epidermal cells, hyp-8 – hyp-11. Hyp-11 is located dorsally, hyp-8 and 9 ventrally, and hyp-10 creates the extreme end of the tail tip; these cells secrete the tail cuticle, with two collagen-rich layers separated by struts (**Fig. 1b**) [9]. The hermaphrodite tail is morphologically fragile because it is thin and consists only of hypodermal cells with no underlying muscle, intestine or gonad tissue to provide additional structural robustness. Indeed, some mutations in genes that regulate membrane lipid homeostasis cause tail end defects in hermaphrodites, suggesting that the Tail End Defects in the hermaphrodite (Ted) phenotype may be a useful indicator of membrane homeostasis defects [10]. Specifically, mutations in *paqr-2* or *iglr-2* primarily result in an excess of saturated fatty acids within membrane phospholipids that in turn causes intolerance to low temperature or dietary saturated fatty acids as well as a Ted phenotype [11, 12]. A previously published screen for mutants that exhibited the same cold intolerance and Ted phenotype as *paqr-2* and *iglr-2* mutants led to the identification of novel mutant allele of *paqr-2* and *iglr-2* as well as of *dpy-23* (involved in endosome recycling to/from the plasma membrane) and *sma-1* (encodes a spectrin that underlies and structurally supports the plasma membrane) [10], adding credence to the possibility that the Ted phenotype may be a useful indicator of membrane homeostasis defects.

**Fig. 1.**
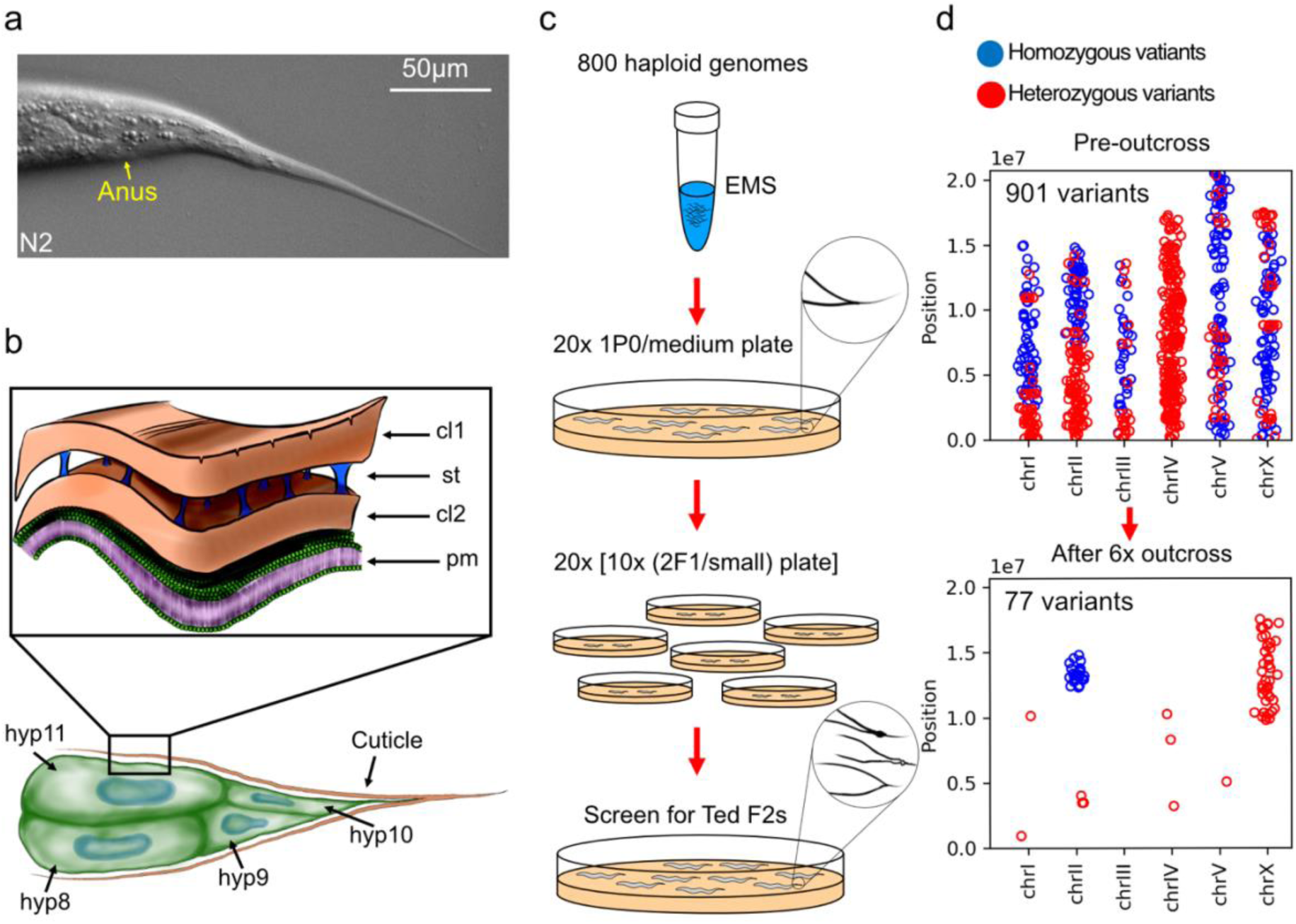
Structure of the tail and overview of the forward genetic screen for Ted (tail end defective) mutants. a) Representative tail of a wild-type hermaphrodite showing a long and tapered tail tip. b) The *C. elegans* hermaphrodite tail is comprised of four epidermal cells (hyp8 – hyp11) that secrete the cuticle through their plasma membrane (pm). The cuticle has two collagen rich layers (cl1 and cl2), separated by struts (st). c) Outline of the forward genetic screen of 800 EMS-mutagenized haploid genomes screened for Ted mutants in the F2 generation. d) The isolated mutants were outcrossed six times and whole genome sequenced to identify clusters of homozygous variants, as shown in this example; the numbers inside the plots represent number of variants found at each step of the outcross.

With the above considerations in mind, we leveraged the Ted phenotype to search for other genetic regulators of cellular boundaries, i.e. cuticle or membrane. We here report the isolation of 21 novel ethyl methanesulfonate (EMS)-induced Ted mutants, the CRISPR/Cas9 verification of 6 of these genes not previously implicated in morphogenesis, and a more detailed study of a novel *ptr-18* allele.

## Results

### Forward genetic screen for Ted mutants

21 Ted mutants were identified by screening the F2 progeny from ∼800 EMS-mutagenized haploid genomes (**Fig. 1c**). After 6 rounds of outcrossing to wild-type worms, genomic DNA was isolated from each mutant and sent for whole genome sequencing. Outcrossing decreases drastically the number of non-specific mutations and helps in identifying clusters of homozygous variants most likely to contain the Ted-causative mutation. An example of pre- and post-outcrossing distribution of heterozygous and homozygous variants is shown in (**Fig. 1d**).

### The screen reveals mutations in regulatory genes

The 21 Ted mutants showed a variety of distinct tail tip morphology phenotypes, indicating that they likely affect different genes and processes. The whole genome sequencing data was plotted on graphs to show the genomic position of all variants with frequency of 0.2 or more and not present in our wild-type parental strain (which was also sequenced) (**Fig. 2a** and **Suppl. Fig. 1**). Assuming recessivity in most cases, we focused on clusters of homozygous GC to AT nucleotide changes (the most common EMS-induced mutation) to identify candidate Ted-causative variants. The Ted mutants were then divided into two groups. The first group contained 6 Ted mutants with candidate mutations in regulatory proteins and that were confirmed by recreating the point mutations using CRISPR/Cas9 (**Fig. 2c and Suppl. Fig. 3**). These novel and confirmed alleles are: *ptr-18(et70), paqr-2(et71), nab-1(et72), ncam-1(et73), vab-9(et74) and efn-4(et75)* (**Fig. 2a-b**). The second group includes Ted mutants that were not CRISPR/Cas9-confirmed because they either contained candidate mutations in cuticular structural proteins or collagen trimmers (13 Ted mutants, including two alleles of *dpy-7* and three of *sqt-3*) or were mutants for which there are no clear candidate mutation (2 Ted mutants) (**Suppl. Fig. 1 and 2**).

**Fig. 2.**
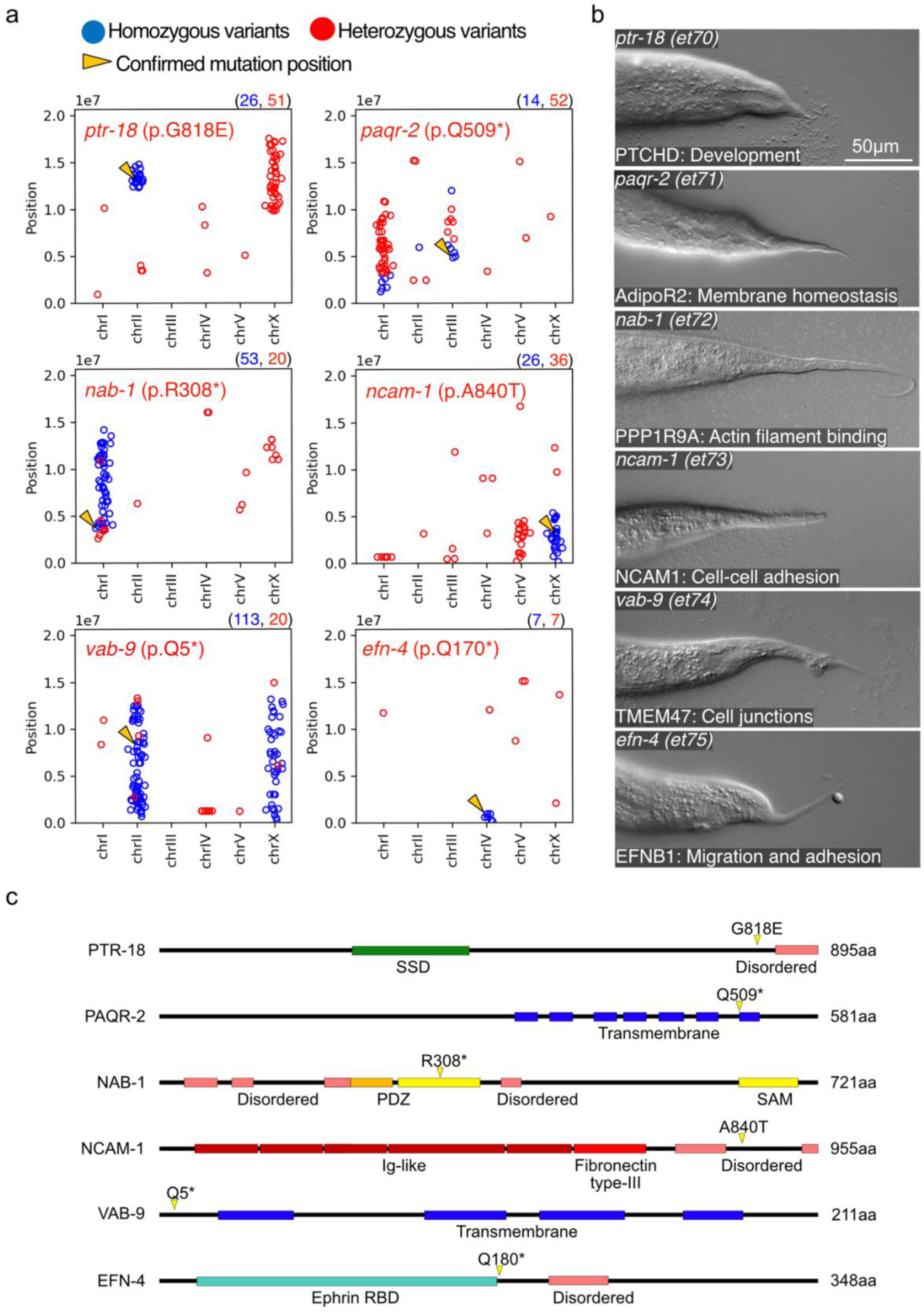
Ted mutants confirmed with CRISPR/Cas9. a) Whole genome sequencing plot showing the position of mutated variant on each chromosome. Red dots represent heterozygous variants; blue dots represent homozygous variants; yellow arrows point to the confirmed mutant alleles positions. The blue and red values above each plot indicate the number of homozygous or heterozygous variants (with frequency ≥ 0.2) present in the sequenced genomes. b) Tails of Ted mutants isolated in the EMS screen with their genotype (upper left corner), and human ortholog name and function (lower left corner). c) Linear models of the proteins encoded by the six CRISPR/Cas9-verified alleles with their functional domains and the mutated amino acid indicated by the yellow arrow.

### Phenotype characterization of Ted mutants

To better understand the cause of the morphological defects in the Ted mutants, we visualized epidermal cells in L3 larvae and adult worms using the adherence junction marker *ajm-1::GFP* (**Fig. 3** and **Suppl. Fig. 4**). All four tail hypodermal cells were present in all the Ted mutants but showed various degrees of constrictions and deformation throughout development/growth. The Ted mutants fell into two groups based on the timing of these morphological defects: a first group with late defects, that initially appear rather normal but in which the Ted phenotype worsens after the L3 stage (including *ptr-18(et70), paqr-2(et71), nab-1(et72), ncam-1(et73))*, and a second group with early defects that show the same phenotype in larval and adult stages (including *vab-9(et74) and efn-4(et75))*. This classification may reflect distinct roles during post-developmental growth (genes in the first group) versus development per se (genes in the second group).

**Fig. 3.**
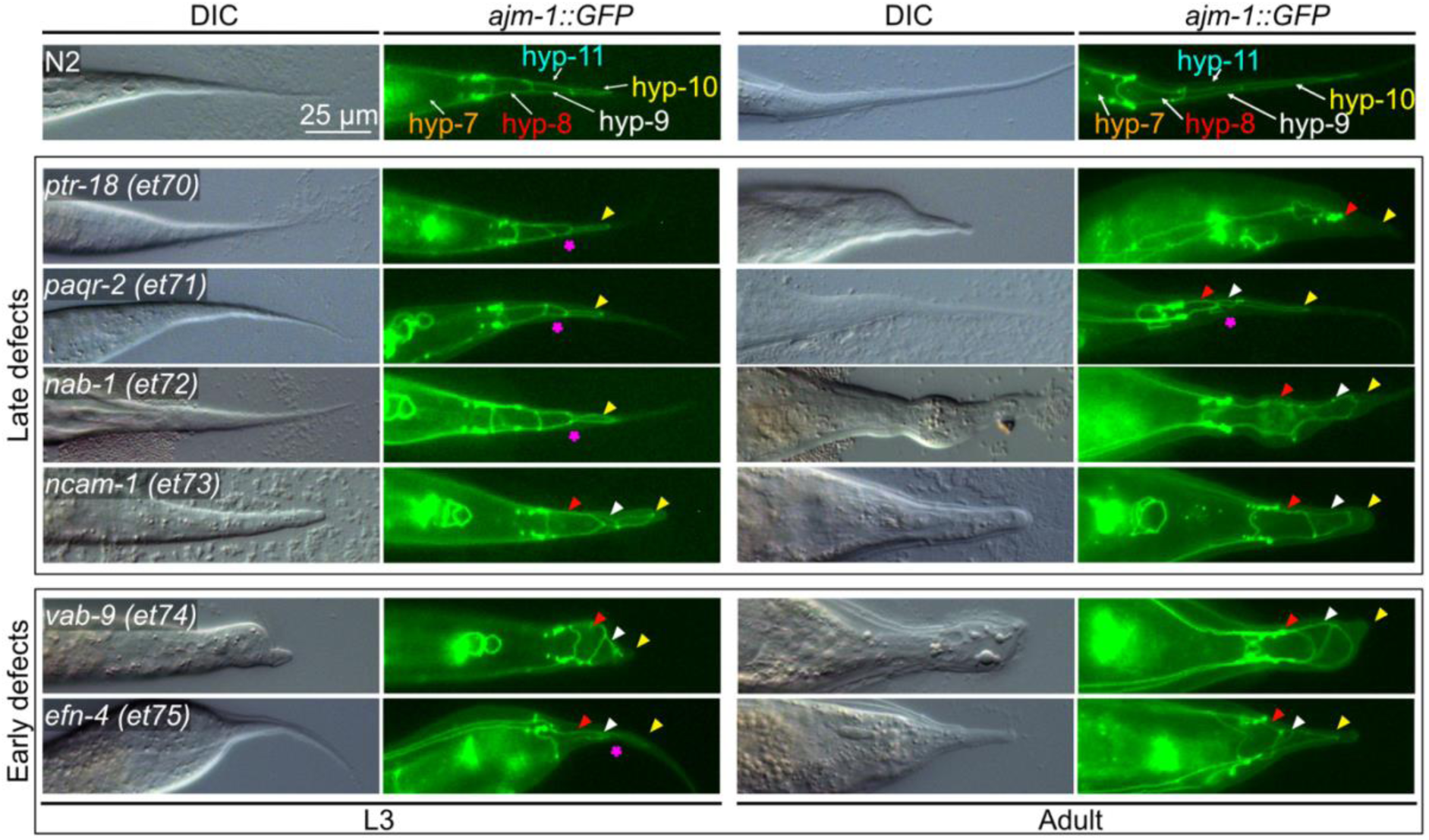
The tail mutants fall into two classes: late defects and early defects. An *ajm-1::GFP* reported was crossed into each Ted mutant to visualize the epidermal cells. Each epidermal cell in the tail is indicated by a color-coded arrowhead (hyp-7: orange; hyp-8: red; hyp-9: white; hyp-10: yellow; hyp-11: cyan). Abnormalities of Ted epidermal cells are pointed out with the appropriately colored arrow. Magenta stars point to abnormal constrictions between hyp-9 and hyp-10.

### Mutation in PTR-18 causes a Ted phenotype

Of the six Ted mutants that were confirmed using CRISPR/Cas9, perhaps the most interesting is the *ptr-18(et70)* allele, which encodes a patch-related (PTR) protein. PTR proteins are evolutionarily conserved and several reports have suggested that an ancestral function for this protein family may be related to membrane homeostasis or lipid transport and thus contribute to ethanol resistance (*ptr-6* [13]), act downstream of *acs-20* (*ptr-8* [14]), or regulate oleic acid metabolism under heat stress (*ptr-23* [15]). The glycine to glutamine substitution at position 818 within the eleventh transmembrane domain (G818E; **Fig. 4a**) in the novel *ptr-18 (et70*) allele or the CRISPR/Cas9 recreated allele *ptr-18(et76)* had nearly the same defective tail phenotype as the *ptr-18(ok3532)* null allele (**Fig. 4b**), suggesting that G818 is a crucial amino acid. The function of *ptr-18* is specific to the hermaphrodite tail since the male tail in *ptr-18(et76);him-5* double mutants is unaffected: the shape of the tail and number of rays is as in wild-type (**Fig. 4c**), and these males were fully fertile. Because some *C. elegans* PTR protein family mutations cause heat stress sensitivity [15], we also characterize the *ptr-18(et76)* at 20°C and 25°C. We measured the length of the worms 48h after L1 synchronization and found that the *ptr-18(et76)* worms had a significantly shorter length at 20°C compared to N2, and that this growth phenotype defect was accentuated at 25°C (**Fig. 4d**). Additionally, while N2 worms uniformly develop within 48h into L4 larvae at 20°C, or adults at 25°C, the *ptr-18(et76)* mutants showed delayed and less uniform development: at 20°C the mutant worms still had a significant percentage of L2 and L3 larvae, while at 25°C L4 worms still accounted for ∼28% of the population (**Fig. 4e**). In conclusion, *ptr-18* is important not only for tail tip morphology but also contributes to developmental rate and robustness, especially at 25°C.

**Fig. 4.**
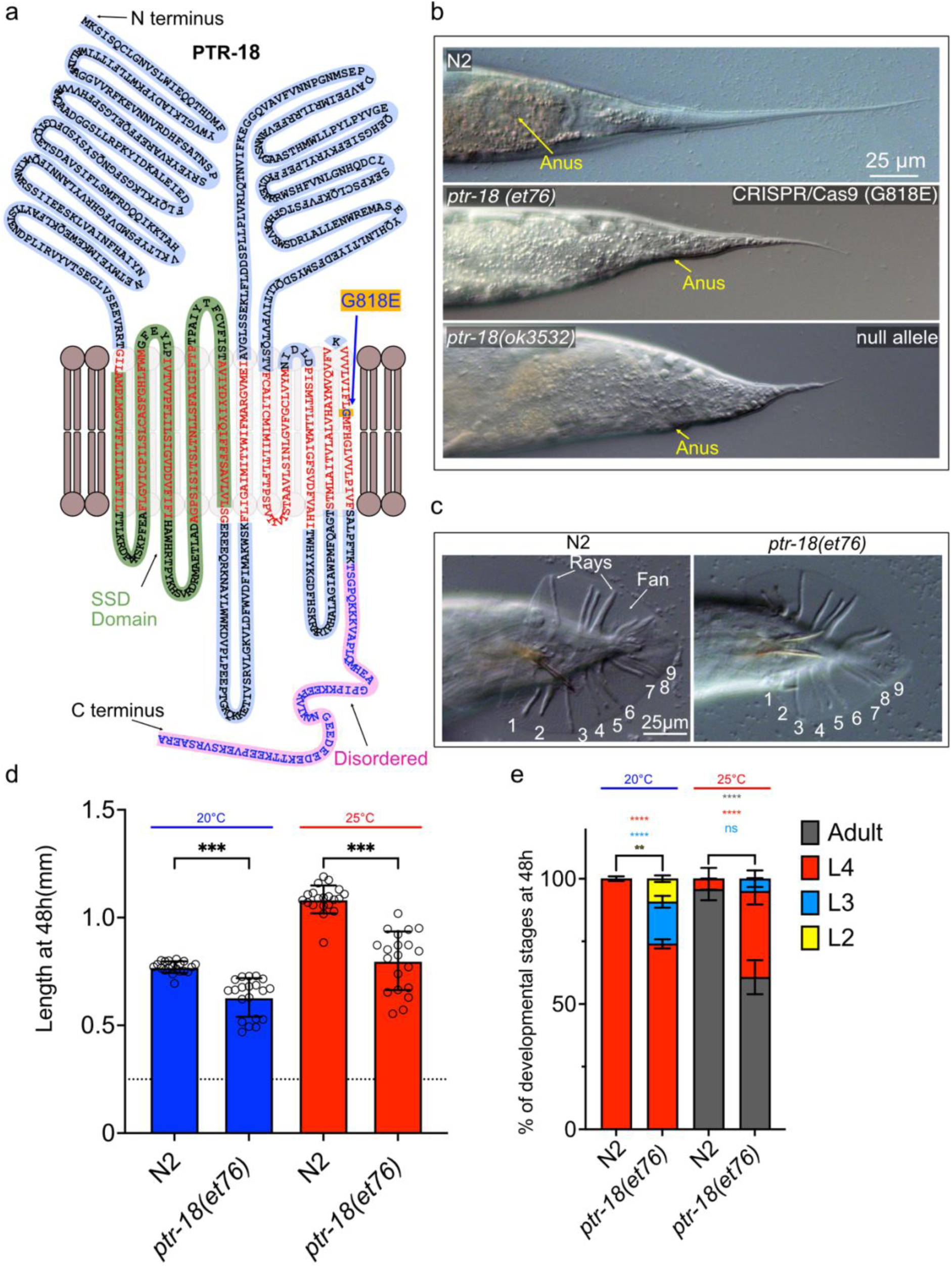
*ptr-18(et76)* is as severe as the null mutant, temperature-sensitive, and does not affect mail tail morphology. a) PTR-18 protein model with the glycine to glutathione amino acid change at position 818 indicated by a blue arrow. b) Comparison between the CRISPR/Cas9-recreated G818E mutation in *ptr-18(et76)* to the *ptr-18(OK3532)* null mutant. c) *ptr-18(et76)* male tail is as in wild-type males. d) Body length comparison at 20°C and 25°C between wild type and *ptr-18(et76)* 48h after synchronization; the dashed line indicates the length of L1 worms at the start of the experiment; n=20 for each strain. Error bars show standard error of the mean. ns p>0.05; *p<0.05, **p<0.01, *** p<0.001, **** p<0.0001 indicate significant difference compared to N2. e) Percentage of larval stages at 20°C and 25°C between wild type and *ptr-18(et76)* 48h after synchronization. n=61 ± 4 from 3 biological replicates. Error bars show standard error of the mean. ns p>0.05; *p<0.05, **p<0.01, *** p<0.001, **** p<0.0001 indicate significant difference compared to N2.

### Localization of PTR-18 is not affected in the mutant strain

We next used CRISPR/Cas9-engineered strains in which the wild-type or mutant *ptr-18* coding sequence was fused to the mNeonGreen reporter to determine whether the G818E amino acid substitution affects the abundance or localization of the PTR-18 protein. We did not observe any significant changes in the protein localization in any of the early developmental stages between N2 and the mutant: the wild-type and mutant PTR-18 proteins localized in epidermal cells and were enriched along the annuli and alae (**Fig. 5a and Suppl. Fig. 5**). However, while the wild-type protein is cleared in later developmental stages, the mutant protein formed persistent aggregates in the tail tips (**Fig. 5a**). To quantify this observation, we incubated L1 larvae at 20°C and 25°C then observed the protein localization once they reached the L4 stage. We found that the G818E substitution caused a temperature-sensitive PTR-18 protein localization phenotype: at 25°C, the wild-type protein is completely cleared in L4 larvae while the mutant protein shows impaired clearance. Additionally, the formation of protein aggregates in the tail tip significantly increased in the mutant strain from 9% at 20°C to 90% at 25°C, and the position of these aggregates correspond to morphological defects visible in DIC images (**Fig. 5b-c**). We conclude that a consequence of the G818E substitution is to impair clearance of the protein once development is complete, leading to the Ted phenotype.

**Fig. 5.**
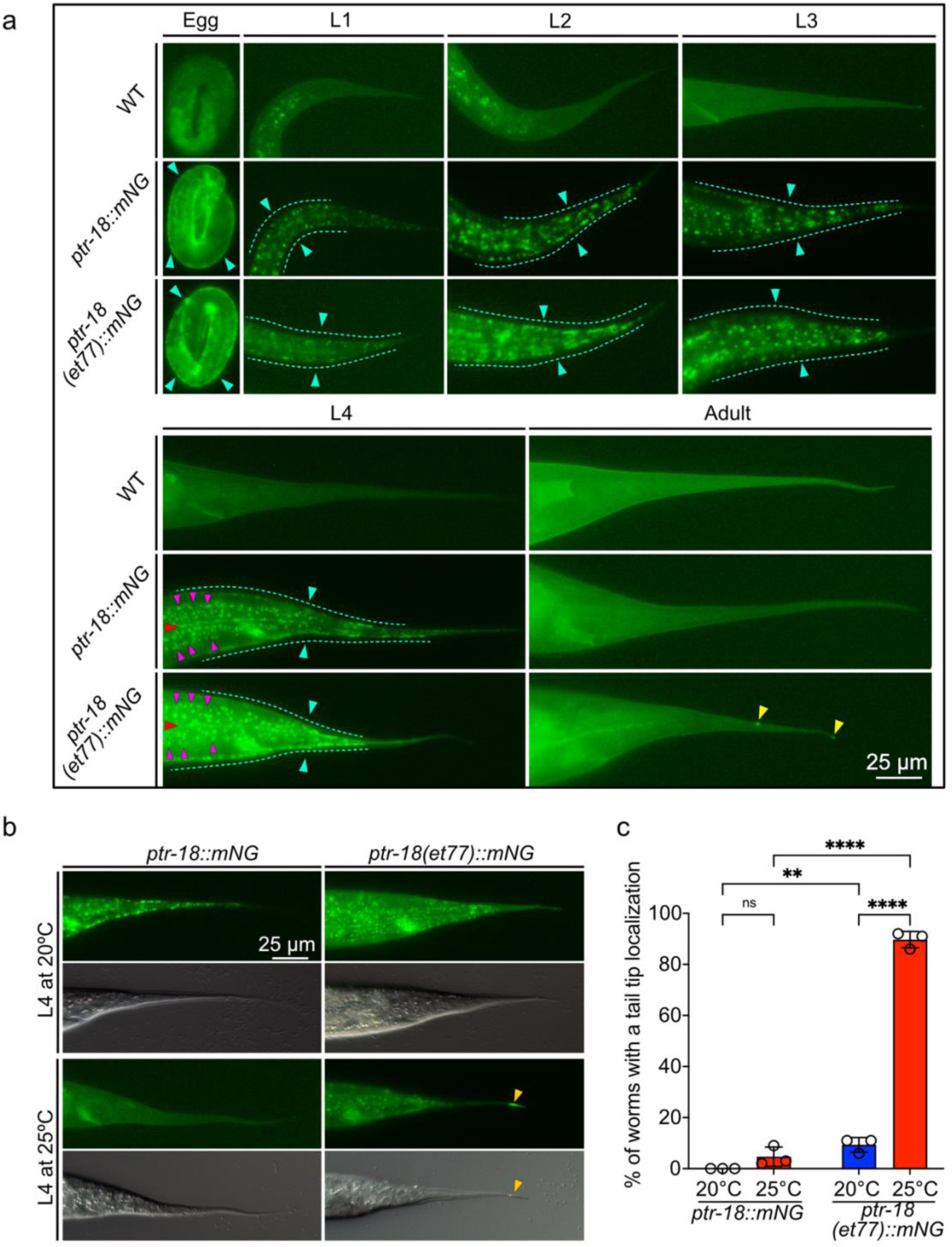
Clearance of the PTR-18(G818E) mutant protein from the tail epidermal cells is impaired during larval development. a) PTR-18 protein localization throughout developmental stages. The PTR-18 protein was labeled with an mNeonGreen tag. Wild type worms were used as a negative control for background autofluorescence. Localization of the protein did not differ between the mutant and wild type worms. Dashed cyan line and arrows are pointing to the distribution of the PTR-18 protein in the epidermis, magenta arrows point to the distribution along the annuli, red arrow points to the distribution along the seam cels, yellow arrow point to PTR-18 aggregates b) The clearance of PTR-18 is increased at 25°C in L4 wild type worms, while the mutant strain has a decreased clearance of PTR-18 and is characterized by the formation of protein aggregates in the tail tip (yellow arrows). c) Percentage of worms with PTR-18 artifacts in the tail tip at 20°C and 25°C between wild type and *ptr-18* mutant worms. Each dot represents a biological replicate with an n=51 ± 14. Error bars show standard error of the mean. ns p>0.05; *p<0.05, **p<0.01, *** p<0.001, **** p<0.0001 indicate significant difference compared to N2.

### The membrane of *ptr-18(et76)* mutants is permeable

Mutants in *paqr-2* [16] and in some members of the PTR protein family [13] have membrane permeability defects. To test whether this is also the case for *ptr-18* mutants, we performed Hoechst staining on synchronized Day 1 adult worms grown at 20°C and 25°C and determined the percentage of worms with stained nuclei in the head or tail region of the worm where auto-fluorescent gut granules do not constitute a confounding factor (**Fig. 6a**). While there was a small, non-significant percentage of N2 worms with permeable membranes at 25°C due to the increased membrane fluidity at higher temperatures, wild type worms were generally non-permeable to Hoechst. For *ptr-18(et76)*, the percentage of worms with permeable membrane varied between experiments, with a mean of 33.67% at 20°C and 51% at 25°C. As a positive control for the test, we used the CRISPR/Cas9-recreated *paqr-2(et77)* of our *paqr-2(et72)* strain, which showed as expected >50% of worms being permeable to Hoechst at 20°C (**Fig. 6b**). We conclude that *ptr-18* is important for establishing impermeable barriers in *C. elegans*, though it is not as crucial for this function as *paqr-2*.

**Fig. 6.**
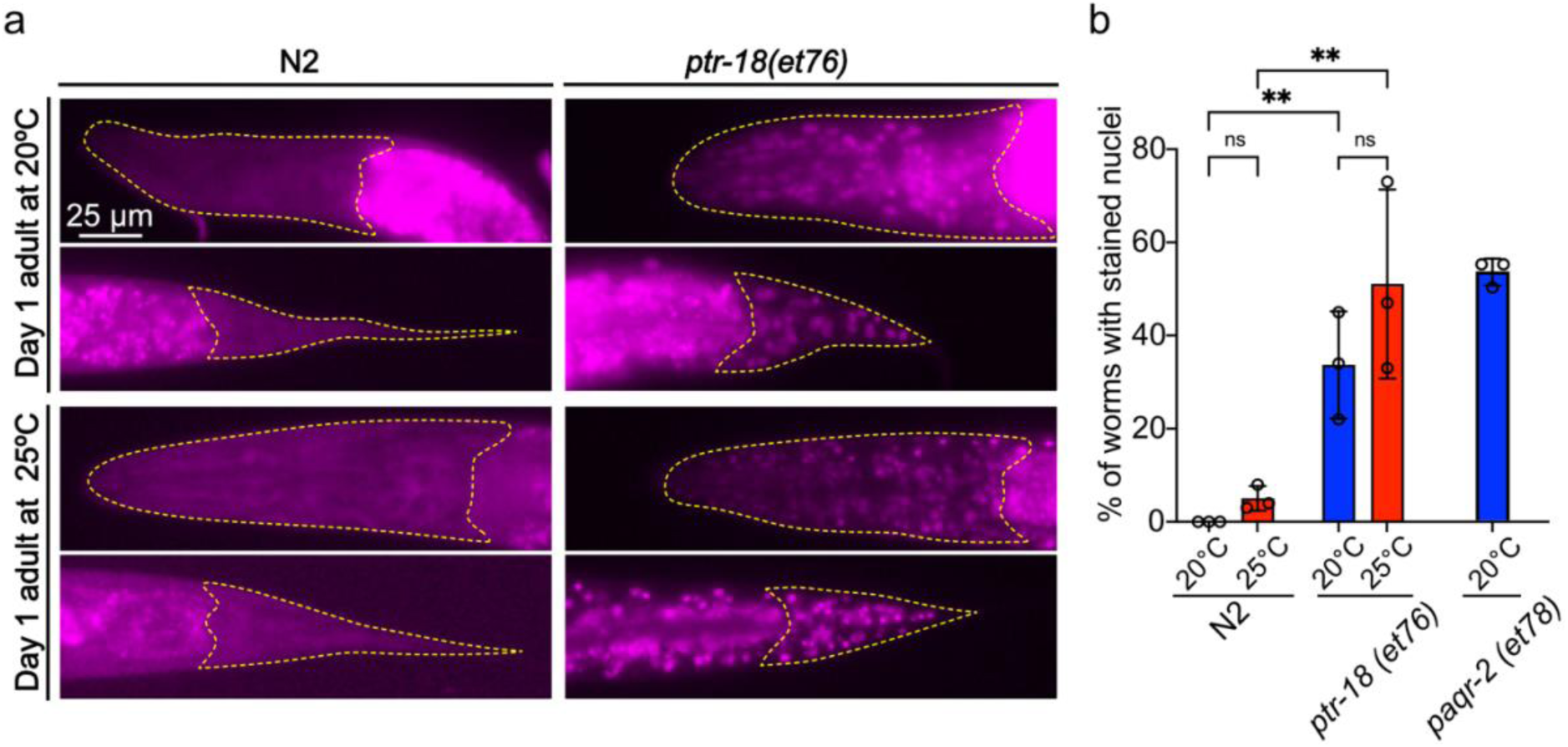
The *ptr-18(et76)* has permeability defects similar to those in *paqr-2* mutants. a) Representative images of wild type and *ptr-18(et76)* worms at 20°C and 25°C with Hoechst 34580 in M9 for 30 minutes. Dashed yellow lines outline the head and tail. b) Quantification of wild type, *ptr-18(et76)* and *paqr-2(et77)* worms with stained nuclei in the head or the tail region at 20°C and 25°C. Each dot represents a biological replicate with an n= >20. Error bars show standard error of the mean. ns p>0.05; *p<0.05, **p<0.01, *** p<0.001, **** p<0.0001 indicate significant difference compared to N2.

### *paqr-2* and *ptr-18* function in separate pathways

As mentioned earlier, the membrane homeostasis *paqr-2(tm3410)* mutant also has a Ted phenotype. To explore the possibility that *paqr-2* and *ptr-18* act in the same pathway, we generated a *ptr-18(et76);paqr-2(tm3410)* double mutant and found that it had a significantly worse phenotype than either single mutant in terms of slowed development, decreased reproduction, and production of many L1-arrested progenies (**Fig. 7a**). We specifically quantified the growth rate of the double mutant and found that it was significantly slower than the single mutants, with a medium length of 0.44 mm after 72h since the synchronization of L1s, i.e. about half the length of either single mutant (**Fig. 7b**). The morphology of the adult worms was also significantly deformed, the tail tip posterior to the anus being completely absent (**Fig. 7c-d**).

**Fig. 7.**
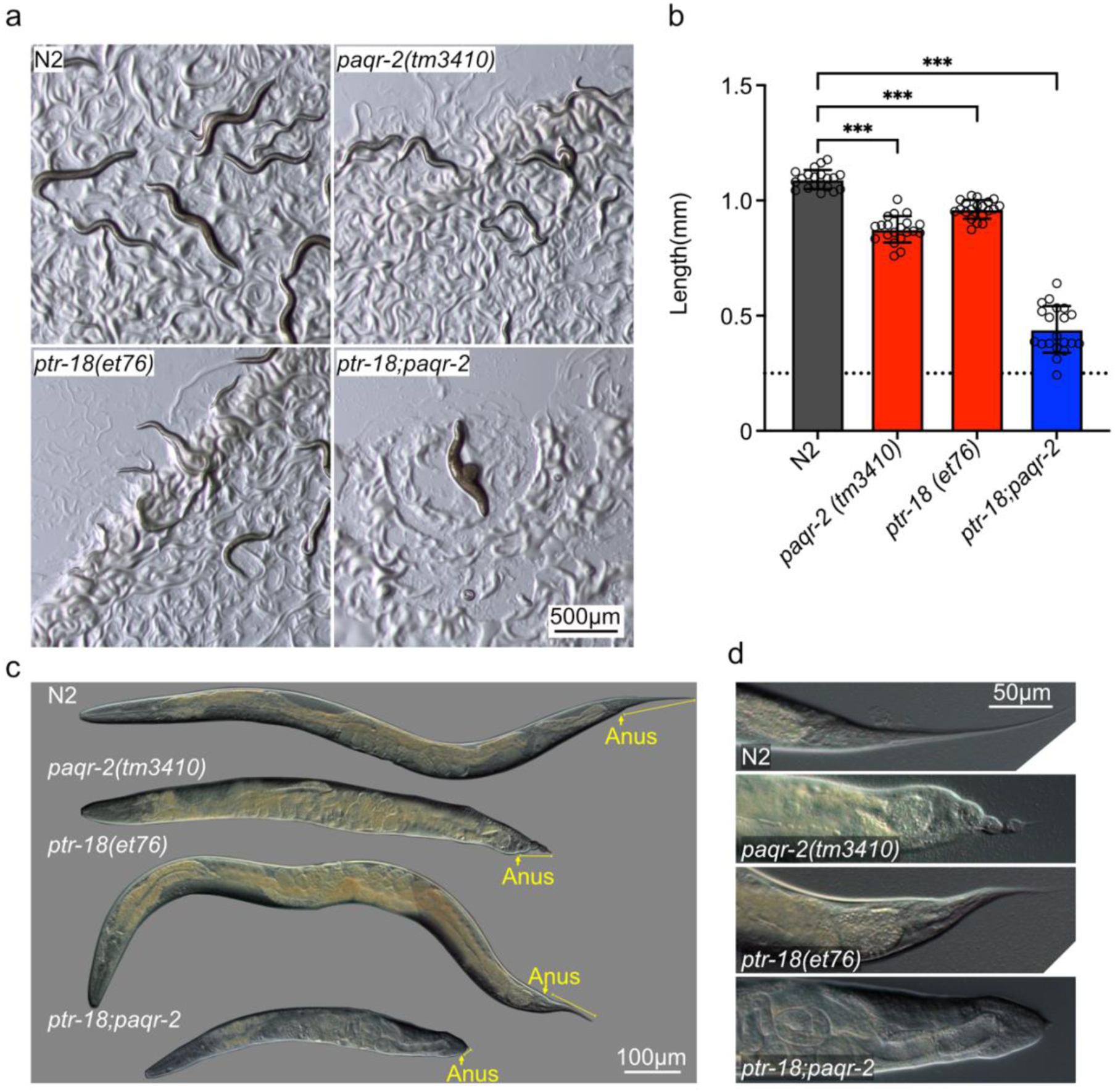
*ptr-18* and *paqr-2* likely act in separate pathways. a) Representative images showing 72 h old plates started with 3 gravid adults for each of the strains. b) Body length measurement 72 h after synchronization; the dashed line indicates the length of L1 worms at the start of the experiment; n=20 for each strain. Error bars show standard error of the mean. *p<0.05, **p<0.01, *** p<0.001 indicate significant difference compared to N2. c) Phenotype comparison of *paqr-2(tm3410)* and *ptr-18(et76)* single mutants, the double mutant and wild type. Yellow arrow points to the anus; yellow line shows the length of the tail. d) Representative images of the tail phenotype for each of the strains.

### Lipid profile of the *ptr-18(et76)* strain is not significantly affected

The tail tip and permeability defects of the *paqr-2* mutant are caused by an excess of saturated fatty acids in its membranes [17–19]. This is not the case for *ptr-18(et76)* even though it too shows tail tip and permeability defects: lipidomics analyses show that the mutant has nearly normal fatty acid composition in phosphatidylcholines and phosphatidylethanolamines, as well as normal overall levels of saturated fatty acids (SFA), monounsaturated fatty acids (MUFA) and polyunsaturated fatty acids (PUFA). Only the fatty acid 19:1 showed a reproducible change, being consistently elevated in the phosphatidylcholines and 15:0 and 20:5 in phosphatidylethanolamines of the *ptr-18(et76)* mutant (**Fig. 8**; a replicate lipidomics profile is shown in **Suppl. Fig. 6**).

**Fig. 8.**
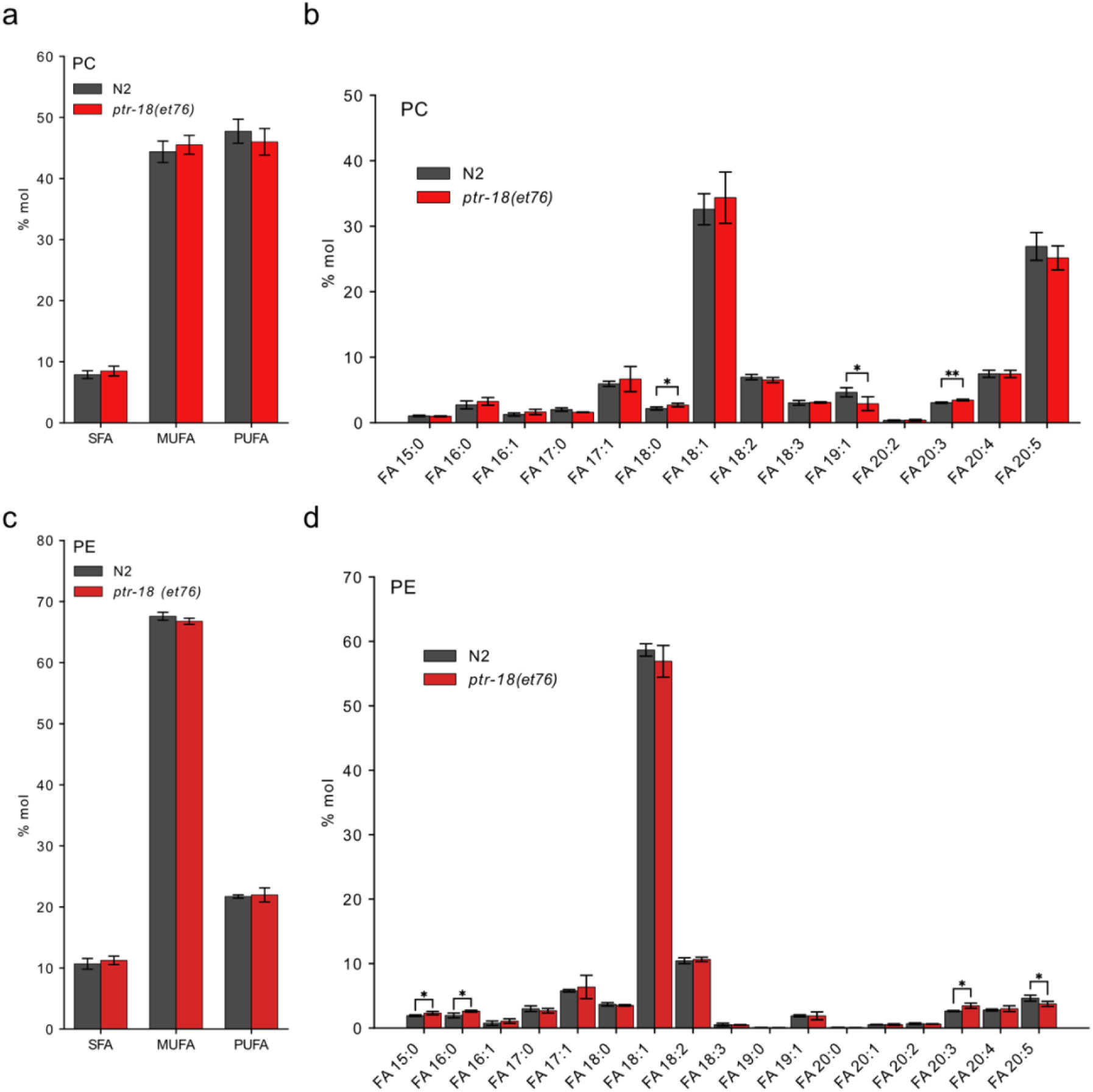
Lipidomics analysis of *ptr-18(et76)* shows no important changes in the lipid composition. a) Levels of SFAs, MUFAs, and PUFAs in PCs between N2 and *ptr-18(et76)* worms are not significantly changed. b) Levels of individual FAs in PCs remain unchanged except three FA species 18:0, 19:1 and 20:3. c) Levels of SFAs, MUFAs, and PUFAs in PEs between N2 and *ptr-18(et76)* worms are not significantly changed. d) Levels of individual FAs in PEs remain unchanged except four FA species 15:0, 16:0 20:3 and 20:5. *p<0.05, **p<0.01, *** p<0.001 indicate significant difference compared to N2.

## Discussion

With the goal to broaden our understanding of the genetic regulation of cellular boundaries, we performed a forward genetic screen of ∼800 haploid genomes and identified 21 mutants with a Ted phenotype. Six of the novel Ted mutants were confirmed using CRISPR/Cas9 and affect genes with diverse functions: cell adhesion (*nab-1, ncam-1*, *vab-9 and efn-4*) [20–22], membrane homeostasis (*paqr-2*) [12] and trafficking/endocytosis (*ptr-18*) [23]. It is worth noting that all either encode membrane-bound proteins, or protein that are functionally adjacent to the membrane, supporting the idea that the Ted phenotype is useful to identify regulators of membranes/cellular boundaries.

Several of the novel Ted mutants affect genes implicated in cell adhesion: NAB-1, EFN-4, NCAM-1 and VAB-9. The novel NAB-1 allele affects an adaptor protein (neurabin) linking transmembrane adhesion molecules to the intracellular active zone proteins and the F-actin network during neuronal polarization of axonal synapses formation [24, 25]. This is reminiscent of a previously identified allele of SMA-1, an actin-associated spectrin, that also causes the Ted phenotype [10]. While NAB-1 functions in a distinct pathway to SMA-1, both are actin filament-binding protein and may affect tail development through related cytoskeletal mechanisms. EFN-4 is one of the ephrin ligands of VAB-1 (EphR) that regulates cell organization and, when mutated, causes morphological defects also in the head and male tail [22]. EFN-4 may also function independently of VAB-1 and acts in the epidermis non-cell autonomously to promote axon guidance and branching [26–28], and we do not know at present if EFN-4 requires VAB-1 for its role in tail tip morphogenesis. NCAM-1, initially characterized as a neural cell adhesion molecule [29], may also contribute to the adhesion between the four tail hypodermal cells in hermaphrodites. As for VAB-9, it is a ubiquitously expressed protein that links the cytoskeleton to adherens junctions and is essential for maintaining F-actin organization and epithelial integrity [21]. Although our *vab-9(et74)* mutant does not show the same phenotype severity as that of the null mutant [21], the early stop codon in the protein is clearly enough to cause developmental problems in the tail. Even though some of the functions of NAB-1, EFN-4, NCAM-1 and VAB-9 are already well established, their role in hermaphrodite tail development was previously unknown.

Our screen for Ted mutants identified a novel *paqr-2* allele. The role of PAQR-2 has been extensively studied in the context of membrane fluidity, where it promotes the desaturation, elongation and incorporation of unsaturated fatty acids into the plasma membrane [16, 30]. Disruption of membrane integrity and of membrane trafficking that contributes to cuticular component secretion in *paqr-2* mutants could be one of main reasons for the characteristic morphological defects. Another of the confirmed Ted mutants is a *ptr-18* allele, which also encodes a protein that likely contributes to the trafficking of secreted protein [23]. The PTR-18 protein is homologous to the human PTCHDs (patched domain-containing proteins). As their names imply, PTCHDs have structural similarities to PTCHs (protein patched homolog); however, despite possessing an SSD (sterol-sensing domain) domain like PTCHs, they do not function in the hedgehog pathway. While the underlying molecular mechanisms of PTCHDs are still not fully resolved, proteins like PTCHD1 regulate neural development and synaptic gene expression [31]. Additionally, in line with its role in neural development, PTCHD1 has been associated with intellectual disabilities and autism [32]. There are 23 PTR proteins in *C. elegans* and they likely have very diverse roles, including lipid metabolism regulation (PTR-8 and PTR-23) and membrane integrity/impermeability (PTR-6) [13–15]. Even though PTR-18 has an SSD domain, like the mammalian patched proteins, and promotes the clearance of the extracellular hedgehog-related protein GRL-7 (groundhog-like) [23], it is important to note that *C. elegans* do not have a hedgehog pathway [33], leaving the exact role of PTR-18 yet to be discerned. Our research shows that an amino acid substitution in the last transmembrane domain of PTR-18 can have a significant effect on the morphology of the worm. Even though the *ptr-18* mutant has permeability defects similar to that of *paqr-2* mutants, the lipidomics data did not reveal any major changes in lipid composition. Others have suggested that defects in the cuticle could indirectly cause membrane permeability defects [5, 34], and this may also be the case for the *ptr-18* mutants. Importantly, while the protein localization during early development was not affected by the mutation, the presence of persistent aggregates in the tail tip points to PTR-18 protein clearance defects between larval stages. This could lead to a buildup of extracellular proteins such as GRL-7, whose localization changes between the matrix and cell during the molting cycle, as others have shown [35].

We did not use CRISPR/Cas9 to confirm 13 of the novel Ted mutants described because they had clear mutations in collagen genes that are structural constituents of the cuticle. Since collagen is the primary component of the cuticle [1], it is unsurprising that a majority (13 of 21, i.e. 62%) of the novel Ted mutants carried mutations in genes that encode or regulate collagen, including two independent alleles of *dpy-7* and three of *sqt-3*. Given that only 800 mutagenized haploid genomes were screened, the sensitivity of Ted phenotype suggests that is especially useful for detecting mutations of the cuticle or extracellular matrix.

Our screen shows that screening for mutants with the Ted phenotype can be a powerful approach to identify novel alleles and pathways that regulate development and cell boundary formation/maintenance. The ease with which mutants were identified and the variety of processes that they affect shows that the four-cell hermaphrodite tail tip is a sensitive structure dependent on many genes for its development (summarized in **Fig. 9**). The variety of pathways that converge on the same Ted phenotype suggests that tail tip morphology and development depend on multi-system integration. Considering that the worm cuticle is a useful model in research concerning wounding, infection, osmotic stress and development [28], our work suggests that the Ted phenotype will be particularly useful to discover new pathways important for these processes.

**Figure 9.**
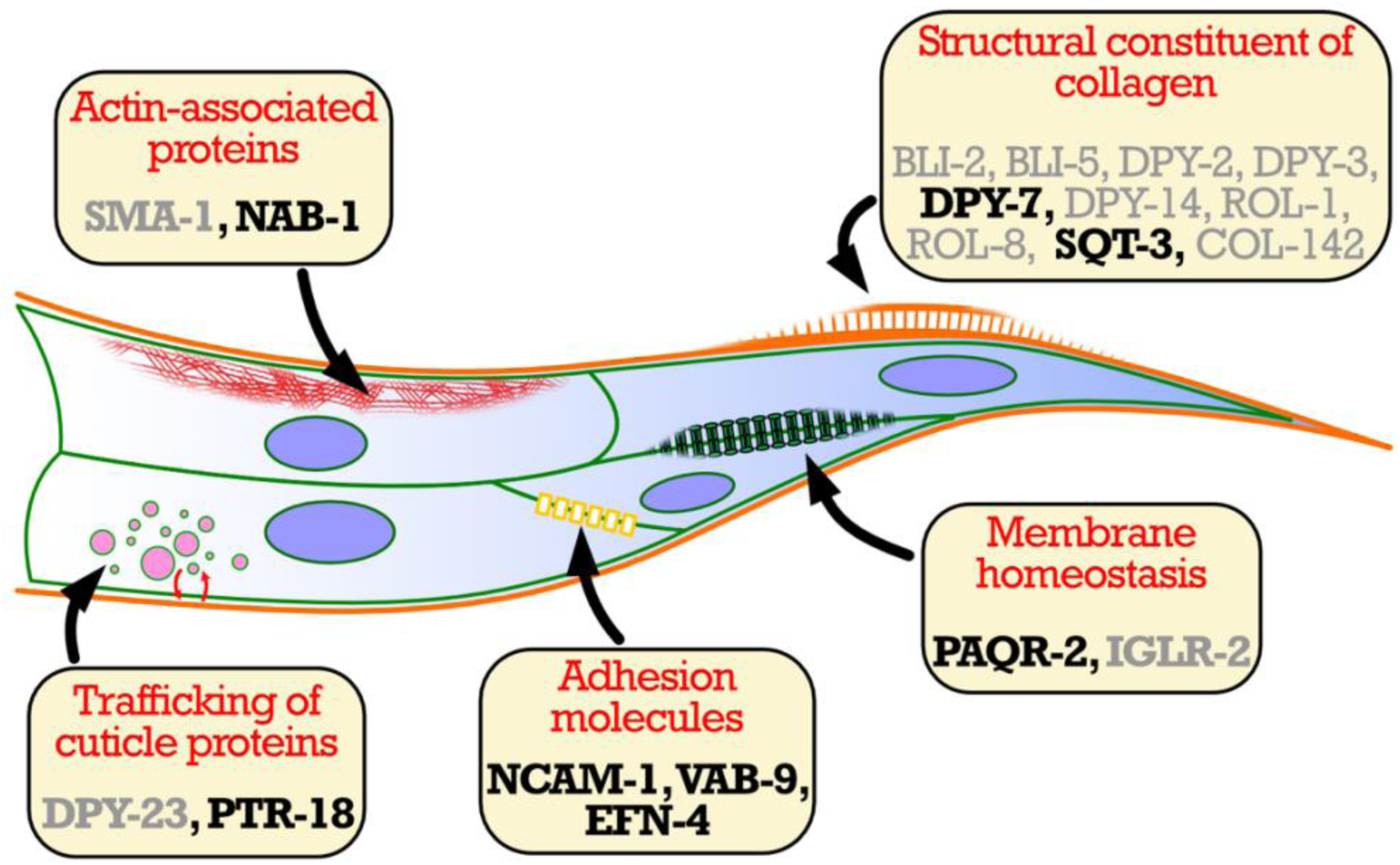
Developmental genetics of the four-cell tail tip. The morphology and development of the *C. elegans* tail tip is dependent on structural constituents of the cuticle and various regulatory protein that include membrane homeostasis, adhesion molecules, actin-associated proteins and trafficking of cuticle proteins.

## Materials and methods

### *C. elegans* strains and cultivation

The wild-type *C. elegans* strain Bristol variety N2, *paqr-2(tm4310), ptr-18(ok3532), jcIs* that carries an *ajm-1::GFP* that expresses the AJM-1::GFP adherens junction fusion protein, and *him-5(e1467)* are available at the *C. elegans* Genetics Center (CGC; USA). The PHX10401 *(ptr-18(syb10401))* strain was created by Suny Biotech using CRISPR/Cas9 and carries the mNeonGreen coding sequence, pDD346, a gift from Daniel Dickinson (Addgene plasmid #133311) was inserted immediately upstream of the *ptr-18* stop codon. The inserted sequence is flanked by 5’- GAAAAATCCGTTCGGTCCGCTGAGCGGGCT -3’ and 5’- aataaaaaattacggaaaacaaaaaagTTA -3’. To make the repaired site resistant to re-cutting, a synonymous mutation (AG to TC) was introduced in the PAM site (AGC).

Unless otherwise stated, worm maintenance and experiments were performed at 20°C, using the *E. coli* strain OP50 as food. The OP50 strain was re-streaked every 6-8 weeks on LB plates and maintained at 4°C. Single colonies were grown overnight at 37°C in an LB medium with constant shaking then spread on NGM plates to produce an *E. coli* lawn after incubation at room temperature. Seeded plates were stored at 8°C for up to ∼2 months.

### Forward genetics screen for Ted mutants

N2 worms were mutagenized for 4h in a 0.05% solution of ethyl methane sulfonate (EMS) according to a standard protocol [36]. After washing, worms were spotted on a NGM plate for 2 h. Among the spotted worms, we singled out 20 L4 hermaphrodites to new plates (progenitor plates) and allowed them to self-fertilize. F1 progeny from each L4 was separated to 10 plates with 2 F1 worms per plate (i.e. four mutagenized haploid genomes per plate) for each of the 20 progenitor plates. Finally, we screened for Ted mutants among the F2 generation. In total, ∼800 haploid genomes were screened. Before detailed characterization, the isolated Ted mutants were outcrossed six times to wild-type worms by three rounds of: mating the Ted mutant with N2 males then mating the resulting heterozygous male progeny with N2 hermaphrodites, then singling out the hermaphrodite progeny and screening for Ted mutants among their offspring.

### Whole genome sequencing

The genomic DNA of the 6 times outcrossed Ted mutants was isolated and sequenced by Eurofins (Constance, Germany) with a mean coverage varying from 36.32X to 81.81 X and their genomes assembled using the 98 Mb *Caenorhabditis elegans* reference genome (Genome assembly WBcel235; NCBI RefSeq assembly GCF_000002985.6). Eurofins applied customized filters to filter out false positive variants using GATK’s Variant Filtration module [37, 38], and the variants detected were annotated based on their gene context using snpEff [39]. For each of the Ted mutants, a hotspot of homozygous variants was identified, and candidate causative variants were selected for further analysis. Of the 21 sequenced mutants, six were experimentally confirmed using CRISP/Cas9.

### CRISPR/Cas9 genome editing

To recreate and confirm the EMS-induced point mutations, we performed CRISPR/Cas9 genome editing as previously described [40, 41]. The genome editing was done utilizing homology-direct repair (HDR) mechanisms. The protospacer-adjacent motif (PAM) site of the ssDNA oligo template was flanked by 40 bp homology arms. Additional to the candidate point mutation, silent mutations were also added around the Cas9 cutting site to prevent re-cutting after editing. Design and synthesis of the ssDNA and CRISPR RNA (crRNA) was done by using the Alt-R HDR Design Tool from IDT (Integrated DNA Technologies, Inc.; Coralville, IA, USA), including proprietary modifications that improve oligo stability. The ssDNA oligos, crRNA sequences and genotyping primers used for CRISPR/Cas9 are listed in Table 1.

The genome editing components were delivered via microinjection into the worm gonads. The injection mix was prepared using 0.5 μl (10 μg/μl) of the Cas9 enzyme (IDT), 5 μl (0.4 μg/μl) tracrRNA (IDT), 2.8 μl (0.4 μg/μl) crRNA (IDT), 2.22 μl (1 μg/μl) of ssDNA (IDT), 40 ng/μl of *pPD118.33* (*Pmyo-2::GFP*), a gift from Andrew Fire (Addgene plasmid #1596), and nuclease-free water to a total volume of 20 μl. Among the F1 generation, worms expressing the *Pmyo-2(GFP)* reporter were isolated and among their progeny we screened for worms with a Ted phenotype. The editing was confirmed by PCR genotyping and successfully edited genes were additionally confirmed by Sanger sequencing (Eurofins).

### Growth assay

For length measurement experiments, bleach-synchronized L1 worms were plated onto NGM plates with OP50 and incubated for 72 h before being mounted and photographed. For experiments performed at 20°C and 25°C the worms were incubated for 48 h. The length was measured using ImageJ.

To determine the distribution of stages at 20°C and 25°C, three replicates of synchronized L1 worms were incubated for 48 h, mounted and photographed. After calculating the percentage of stage distribution for each replicate, we plotted the mean distribution of the replicates.

### Hoechst staining

Bleach-synchronized L1 worms were incubated at 20°C or 25°C and Day 1 adults were washed with M9 then incubated for 30 minutes at room temperature in a 1 μg/ml solution of Hoechst 34580 dissolved in M9. After staining, the worms were washed twice with M9, mounted on agarose pads and imaged using a Zeiss Axioscope microscope. Anterior pharyngeal and posterior tail regions were used for quantification to avoid interference from intestinal gut granule autofluorescence.

### Lipidomics

For each strain, four independent replicates of synchronized L4 worms were grown on 9 cm diameter NGM plates; this was done for each of two separate experiments. The worms were collected by washing 3 times with M9, pelleted with as much as possible of the supernatant removed and stored at -80°C until analysis [42]. For lipid extraction, the pellet was sonicated for 10 minutes in methanol;butanol [1:3] and then extracted according to published methods [43]. Lipid extracts were evaporated and reconstituted in chloroform:methanol [1:2] with 5 mM ammonium acetate. This solution was infused directly (shotgun approach) into a QTRAP 5500 mass spectrometer (Sciex) equipped with a TriVersa NanoMate (Advion Bioscience) as described previously. Phospholipids were measured using precursor ion scanning in negative mode using the fatty acids as fragments [44, 45]. To generate the phospholipid composition (as mol%) the signals from individual phospholipids (area under the m/z peak in the spectra) were divided by the signal from all detected phospholipids of the same class. The data were evaluated using the LipidView software (Sciex).

### Statistics

Error bars show standard deviations unless otherwise stated. The statistical analysis for worm length measurement was done in GraphPad Prism with a one-way ANOVA test to identify statistical significance from the N2 control. For the distribution of worm stages, permeability assay, and PTR-18 protein distribution, a two-way ANOVA test was done. The statistical analysis of the lipidomics data was analyzed with a student *t-*test in Pythons 3.0 with NumPy, comparing the values between N2 and *ptr-18(et76)* for each FA individually. The plotting of the graphs was done with seaborn and matplotlib [46–48]. Every experiment was repeated at least twice, and the statistical analysis shown applies to the presented experimental results.

## Data availability

The authors affirm that all data necessary for confirming the conclusions of the article are present within the article, figures, and tables. Strains and plasmids are available upon request.

## Acknowledgements

This work was supported by the Swedish Research Council grants 2020-03300, Cancerfonden grant 19 0029, and Wilhelm och Martina Lundgrens Vetenskapsfond grant 2025-GU-5010. Some strains were provided by the CGC, which is funded by NIH Office of Research Infrastructure Programs (P40 OD010440).

